# Phase dynamics of cerebral blood flow in subarachnoid haemorrhage in response to sodium nitrite infusion

**DOI:** 10.1101/2020.06.15.152108

**Authors:** Martyn Ezra, Payashi Garry, Matthew J Rowland, Georgios D Mitsis, Kyle TS Pattinson

**Affiliations:** Nuffield Division of Anaesthetics, Nuffield Department of Clinical Neurosciences, University of Oxford, UK; Department of Bioengineering, McGill University, Canada

**Keywords:** Nitrite, Subarachnoid Haemorrhage, Stroke, Nitric Oxide, Cerebral Blood Flow, Cerebral Autoregulation

## Abstract

Aneurysmal subarachnoid haemorrhage (SAH) is a devastating subset of stroke. One of the major determinates of morbidity is the development of delayed cerebral ischemia (DCI). Disruption of the nitric oxide (NO) pathway and consequently the control of cerebral blood flow (CBF), known as cerebral autoregulation, is believed to play a role in its pathophysiology. Through the pharmacological manipulation of *in vivo* NO levels using an exogenous NO donor we sought to explore this relationship.

Phase synchronisation index (PSI), an expression of the interdependence between CBF and arterial blood pressure (ABP) and thus cerebral autoregulation, was calculated before and during sodium nitrite administration in 10 high-grade SAH patients acutely post-rupture. In patients that did not develop DCI, there was a significant increase in PSI around 0.1 Hz during the administration of sodium nitrite (33%; p-value 0.006). In patients that developed DCI, PSI did not change significantly.

Synchronisation between ABP and CBF at 0.1 Hz has been proposed as a mechanism by which organ perfusion is maintained, during periods of physiological stress. These findings suggest that functional NO depletion plays a role in impaired cerebral autoregulation following SAH, but the development of DCI may have a distinct pathophysiological aetiology.

## 1 Introduction

Aneurysmal subarachnoid haemorrhage (SAH) is a devastating subset of stroke, characterised by high patient mortality and long-term disability.[1] As a result, SAH is estimated to cost the UK economy £510 million/year[2], with the loss of productive years equal to that of ischaemic stroke. One of the major causes of morbidity and mortality is the development of secondary neurological injuries, 3-14 days after the initial bleed, previously termed “vasospasm” but now known as delayed cerebral ischemia (DCI).[3] The underlying aetiology of DCI is poorly understood, despite it affecting 30% of SAH patients who survive to hospital admission.[4] The phenomenon of early brain injury (EBI) has recently attracted attention as an important mechanism in the development of DCI. The term refers to a multifactorial global cerebral injury that develops in the first 72 hours following SAH.[5]

A growing body of clinical studies suggest that disruption to cerebral blood flow (CBF) control may play a critical role in the pathophysiology of EBI, and that its severity directly correlates with the development of DCI.[6–14] As a consequence, the role of the nitric oxide (NO) signalling pathway EBI has garnered considerable interest.[15]

Animal models suggest that NO levels demonstrate a biphasic response following SAH. Haemoglobin and its breakdown products, buffer free NO and inhibit endothelial NO synthase (eNOS) function resulting in profound early nitric oxide depletion. This followed at 6-72 hours by an increase in inducible NO synthase (iNOS) activity and an elevation of NO above baseline levels.[16,17] Whilst studies in humans almost universally demonstrate elevated NO products in the EBI period, evidence corelating changes in NO and the develop of DCI is far from conclusive. Both relatively decreased [18,19] and increased [20,21] levels have been shown to demonstrate an association with the development of DCI.

One of the primary obstacles in determining the role of NO is the inability to measure NO levels directly due to its exceptional short biological half-life. NO levels are extrapolated from the measurement of NO breakdown products within the cerebrospinal fluid (CSF), and therefore, neither account for regional distribution or biological availability of NO. Alternatively, it may be possible to probe the NO pathway by determining the cerebral response to *in vivo* manipulation of the NO levels using an exogenous nitric oxide donor.

During the last decade, a series of pre-clinical and clinical studies have demonstrated that the inorganic anion nitrite represents an important endogenous storage pool of NO.[22] In ischemic or hypoxic tissue environments, where enzymatic production of NO is limited by the requirement for molecular oxygen, nitrite is reduced to form NO. Critically the magnitude of this storage pool is modifiable by loading with exogenously administered nitrite.[23]

Following the administration of sodium nitrite, Garry et al., demonstrated significant increase in the electroencephalographic (EEG) spectral alpha/delta power ratio (ADR) in high grade SAH patients.[24] ADR has been shown to correlate with regional CBF after ischemic stroke. [25] Furthermore, primate autologous blood clot injection models of SAH suggest that sodium nitrite may prevent or reverse the cerebrovascular manifestations of DCI. [26,27] These findings support the hypothesis that disruption to the NO pathway plays an important role in the disruption of CBF control during EBI. However, direct examination of the relationship between the NO pathway, CBF control and EBI has not previously been investigated.

The control of CBF with respect to arterial blood pressure (ABP) is classical referred to as cerebral autoregulation.[28] Utilising induced changes or naturally occurring spontaneous oscillations in ABP it is possible to characterise the dynamic relationship between ABP and CBF mathematically and, therefore, to derive metrics to quantify dynamic cerebral autoregulation (dCA).[29] Normal dCA is often described in terms of a high pass filter, where low frequency changes in ABP are dampened.[29]

Efforts to demonstrate that disruption to the NO signalling pathway has a discernible effect on dCA utilising healthy volunteer models have been so far inconclusive.[30–32] This may be in part due to the methods chosen to determine dCA, which assume that dCA can be described as a linear time-invariant system. However, both these assumptions are likely to be untrue, especially at low frequencies where non-linearity and non-stationarity have a pronounced effect.[33–35]

To address non-stationarity, Latka et al. utilised the continuous wavelet transform[36], an integral transform that uses wavelets, which are essentially wave-like functions with compact support i.e. they are zero outside some finite time interval (Figure 1a).[37] Wavelets are localised in both time and frequency; thus, they can be used to extract the time-varying spectral features of a signal. By determining the instantaneous phase of the ABP and CBF signals, Latka et al. derived a statistic, the phase synchronisation index (PSI) (γ), to describe the interdependence of the two signals in relation to the variance of their phase difference over time.[36] Repeated studies have shown that the PSI of healthy volunteers demonstrates a highly consistent morphology (Figure 1b).[38–41]

**Figure 1:**
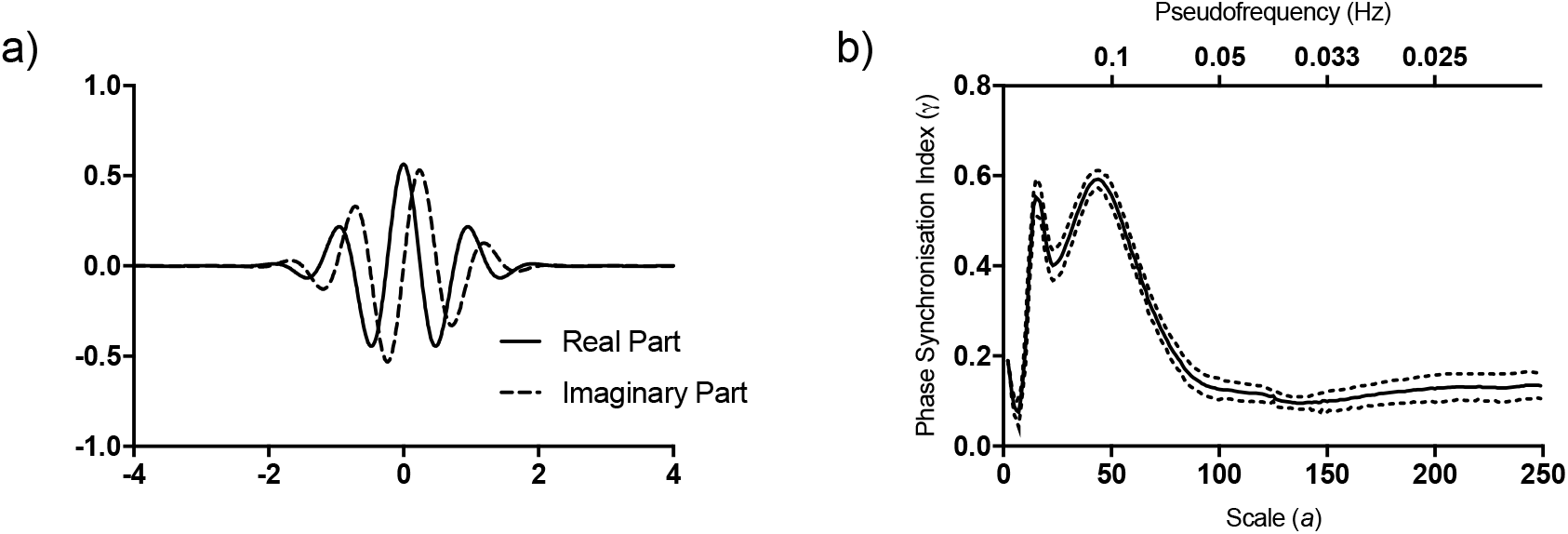
a) Real (solid line) and Imaginary (dashed line) parts of the complex mother Morlet wavelet. b) Phase Synchronization Index (*γ*) as a function of scale a (bottom x-axis) or pseudofrequency *f*_*a*_ (top x-axis) in healthy volunteers adapted from Latka et al. Solid line represents the group average, dashed lines correspond to the standard error (SE). Figure demonstrates the characteristic peaks at scale 15 (0.33 Hz) and scale 50 (0.1 Hz), as well as low phase synchronisation at scales > 75 (<0.07 Hz). These were proposed by Latka et al. to represent respiration rhythms, Mayer waves and intact autoregulation, respectively[36].

We propose that examination of changes to the characteristic morphology of the PSI may represent a more sensitive method of determining the effects of NO on dCA. As a result, the aim of the study was to examine the role of the NO pathway on dCA during EBI in high grade SAH using pharmacological manipulation of *in vivo* NO levels and utilising a novel non-stationary analysis technique. We hypothesised that SAH would be associated with impaired CBF phase dynamics, and that manipulating the NO pathway with intravenous sodium nitrite would result in normalisation of this relationship.

## 2 Methods

### 2.1 Participants

In this study we performed an extended analysis of the haemodynamic data collected by Garry et al.[24] Of the 14 patients included in the original study, 10 patients had continuous transcranial Doppler ultrasonography (TCD) and ABP waveforms recorded of sufficient quality required for the present analysis.

All patients aged 18-80 years admitted to the John Radcliffe Hospital, Oxford after having suffered severe aneurysmal SAH (World Federation of Neurosurgeons (WFNS) grade 3-5) and successful treatment with endovascular coiling were eligible for inclusion. Exclusion criteria included contraindications to sodium nitrite, specifically, severe cardiovascular compromise and pre-existing methaemoglobinaemia. Written informed consent was obtained from the next of kin, and from participants themselves, if they regained capacity to consent. The study was approved by the South Central-Oxford C NHS Health Research Authority Ethics Committee 12/SC/0366.

All patients were managed in line with local clinical guidelines on the intensive care unit. Sedation was maintained with a combination of propofol, fentanyl, and midazolam with neuromuscular blockade using atracurium as required. Mean arterial blood pressure (MAP) was maintained between 60-90 mmHg using an infusion of noradrenaline. All patients received oral nimodipine 60mg 4 hourly and had plasma magnesium levels maintained at 1-1.2 mmol/L. Patients with hydrocephalus were treated with external ventricular drainage.

### 2.2 Study Design and Data Collection

All participants received an infusion of sodium nitrite at 10 mcg/kg/min for one hour via a peripherally sited intravenous canula. Sodium nitrite has been established as safe for long term use in human SAH.[42] In this instance, the dosing schedule was developed as a compromise between ensuring adequate plasma nitrite levels and minimisation of cardiovascular effects.

Continuous measurements of middle cerebral artery (MCA) blood flow velocity, arterial blood pressure (ABP), intracranial pressure (ICP) and end tidal CO_2_ (P_ET_CO_2_) and O_2_ (P_ET_O_2_) were made prior to and during the infusion of sodium nitrite.

CBF velocity (CBFV; cm/s) in the MCA was monitored using TCD (EZ-Dop, DWL, Germany; 2 MHz). Insonation of the MCA M1 segment was performed using a probe fixed at a constant angle and position relative to the head with a custom-made headset. This was initially performed bilaterally to assess for the best temporal window and then continuously recorded unilaterally from the best side.

ABP (mmHg) was monitored via an intraarterial catheter sited in the radial artery (Baxter Healthcare, UK). ICP (mmHg) was monitored with an intraparenchymal microsensor inserted into the frontal lobe (Codman, UK). 3 patients did not have ICP monitoring. End tP_ET_CO_2_ and P_ET_O_2_ (kPa) were monitored using side stream infrared and optical sensors, respectively (AD Instruments, New Zealand). All data was sampled at 100 Hz on a Power-1401 data acquisition interface (Cambridge Electronic Design, UK) and recorded for off-line analysis.

All drug infusions were documented and changes in rate were minimised for the duration of the recording. The diagnosis of delayed cerebral ischemia/infarction was made by the attending clinicians, independent to the study, in line with consensus multi-disciplinary guidelines.[43] We have explicitly avoided the use of the term “vasospasm” to avoid confusion between angiographic vasospasm and the clinical manifestations of cerebral ischemia which historically has also been termed vasospasm. Each surviving patient was followed up at 3-6 months post-rupture using the modified Rankin scale (mRS).[44]

### 2.3 Data Pre-processing

Data pre-processing was carried out using custom written MATLAB[45] scripts. Baseline data consisted of 30 minutes of recordings made prior to the infusion. Infusion data consisted of 45 minutes of recordings made 5 minutes after the start of the infusion, to account for dead space in the peripheral canula and connectors.

Data was divided into 15 minute segments, in line with Latka et al.[36] and visually screened to exclude those containing excessive artefacts lasting longer than three cardiac cycles. Short periods of artefact, less than three cardiac cycles, were removed and replaced by linear interpolation.[46] This resulted in the exclusion of 4 out of 50 data segments (8% of data).

Beat-to-beat average values of CBFV, ABP and ICP were obtained via waveform integration using the time of the BP diastolic value to indicate each cardiac cycle. The non-uniformly spaced timeseries were resampled uniformly at 5 Hz using spline (third-order polynomials) interpolation. No detrending, normalization, or filtering was performed, in line with the CARNET (Cerebral Autoregulation Network) transfer function analysis white paper.[47] P_ET_CO_2_ values were calculated via waveform peak identification.

### 2.4 Wavelet Phase Synchronisation

A continuous wavelet transform was performed on all ABP and CBFV waveform segments using the MATLAB CWT function.[45] Continuous wavelet analysis is performed by starting with a prototype wavelet, known as the mother wavelet (*ψ*), that is scaled (i.e dilated or shrunk) in such a way that the wavelet shape remains constant, but the duration is altered. This results in the formation of a daughter wavelet at each scale (*a*). The daughter wavelet is then translated along the input signal and convolved with the signal at each time point. Varying the wavelet scale and translating along the localised time index, enables the resolution of both the amplitude of any features, versus the scale, and how this amplitude varies with time.

The continuous wavelet transform of a signal *s*(*t*), for all real positive values of scale *a* and time delay *t*_0_ is given by:

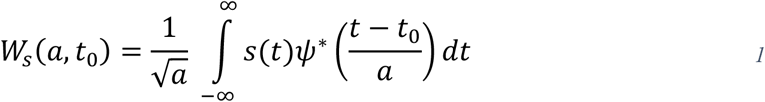

Where 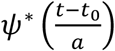 denotes the scaled and translated complex conjugate of the mother wavelet (ψ), 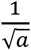 represents the scaling factor at each scale and *W*_*s*_(*a*, *t*_0_) the wavelet coefficients.

A relationship between the scale of the daughter wavelet and frequency can then be approximated using:

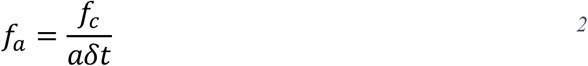

Where *f*_*a*_ is the pseudofrequency of scale *a* and *f*_c_ is the central frequency of the mother wavelet. For comparability with the majority of studies investigating CBF control, we will primarily refer to frequency rather than scale.

Input signals are finite in length, however, the wavelet function extends beyond the borders of these timeseries. As a consequence, the resulting convolution will exhibit edge effects. It is therefore necessary to define a cone of influence (COI), for which edge effects cannot be ignored and are therefore excluded from the analysis. We consider the COI as the area that the wavelet power has dropped to *e*^−2^ of the value at the edge.[48] The COI for larger scales (where the daughter wavelet is dilated to a greater degree) will occupy a larger proportion of the timeseries. The larger the scale, the longer the daughter wavelet and, therefore, extension beyond the border of the timeseries.

It possible to resolve the amplitude of a signal by time and scale and also by the phase. By utilising a complex mother wavelet, i.e. one that contains both real and imaginary components, the instantaneous phase of the signal (*ϕ*_*s*_) at a given scale *a* is given by:

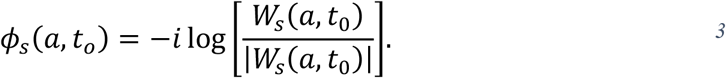

where |*W*_*s*_(*a*, *t*_0_)| is the compex modulus of the complex wavelet coefficients.

In this analysis, we employed the complex Morlet wavelet with bandwidth *f*_b_ and centre frequency *f*_c_, both equal to 1, as per Latka et al.[36] The equation defining the complex mother Morlet wavelet is given by:

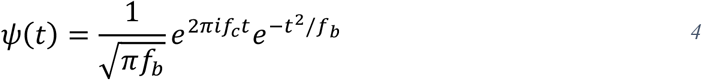

The real and imaginary parts of the complex Morlet are plotted in Figure 1a. As can be seen, the Morlet wavelet is a sinusoidal wave modulated by a Gaussian function.

From the instantaneous phase representations of the ABP and CBFV timeseries, *ϕ*_*P*_(*a*, *t*_0_) and *ϕ*_*V*_(*a*, *t*_0_), respectively, the instantaneous phase difference Δ*ϕ*(*a*, *t*_0_) can be defined as:

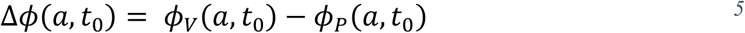

Examination of the distribution of phase difference can then be used to characterise the synchronisation of the two timeseries. A uniform distribution would represent absent synchronisation, i.e. the two timeseries are statistically independent. On the other hand, two timeseries which are synchronised (phase locked) would have a sharp peak in the distribution of phase difference i.e. Δ*ϕ* ≈ constant. Latka et al., quantified the stability of the time-resolved phase difference with an inverse circular statistical analogue of variance using:

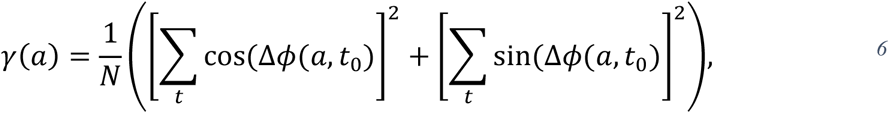

where *γ*(*a*) represents the phase synchronisation index (PSI) at scale *a* that lies in the interval 0 ≤ *γ* ≤ 1, and *N* is equal to the number of time points. A PSI (*γ*) of 1 represents perfectly synchronised timeseries at a given scale. On the other hand, when the distribution of the phase differences is uniform, i.e. perfectly independent, the time average of both trigonometric functions are zero, which leads to a PSI (*γ*) of 0.

Figure 2a demonstrates the normalised phase difference (Δ*ϕ*/π) against time at each pseudofrequency/scale (*HZ*/*a*) in a single patient. Figure 2b demonstrates the derived PSI (*γ*) in the same patient for values outside the COI. At 0.33 Hz (scale 15), the PSI (*γ*) is high. If we examine the instantaneous phase difference plot at this frequency (Figure 2c), we can see that the phase difference varies very little over time, as manifested by the narrow distribution in the corresponding histogram plot (Figure 2d). In contrast, at 0.05 Hz (scale 100), where the PSI (*γ*) is very low, the phase difference varies greatly over time (Figure 2e), with a broad distribution (Figure 2f).

**Figure 2:**
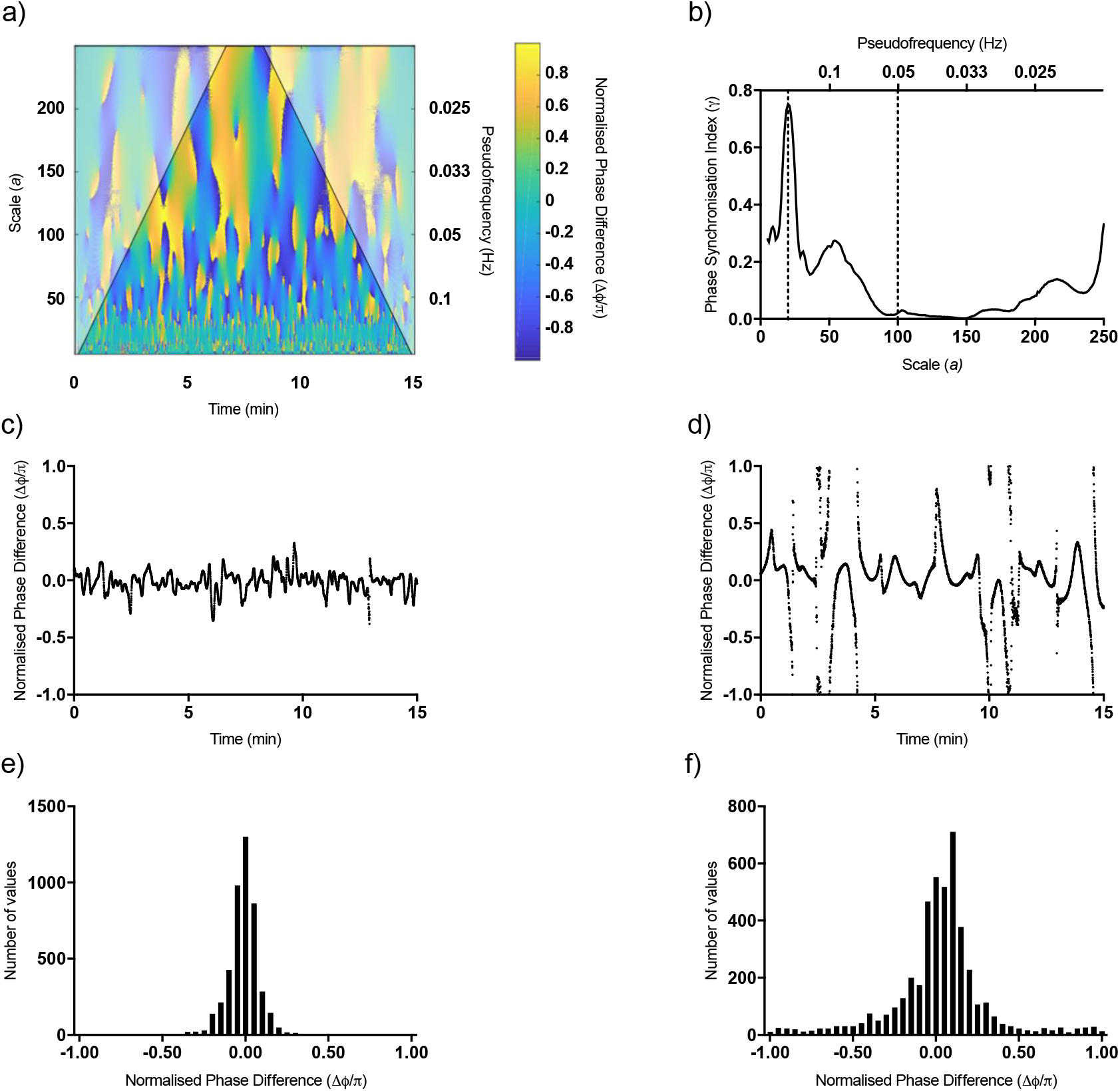
Instantaneous phase difference for a representative subject. a) Time-scale/frequency density map of the normalised instantaneous phase difference (Δ*ϕ*/*π*). The shaded represents COI. b) Plot of the Phase Synchronisation Index (*γ*), demonstrating high phase synchronisation at 0.33 Hz (scale 15) and low phase synchronisation at 0.05 Hz (scale 100) (dotted lines). Normalised phase difference over time at c) 0.33 Hz demonstrates a relatively constant phase difference overtime, whereas at d) 0.05 Hz large variability in phase difference over time can be observed. Histogram of phase difference at e) 0.33 Hz and f) 0.05 Hz, demonstrating small and large variance respectively

### 2.5 Statistical Analysis

Statistical analysis of the PSI was carried out using R (R Foundation for Statistical Computing)[49] and the analysis package nlme (R package version 3.1-131).[50] To determine the effect of sodium nitrite administration and interaction with the development of DCI, a two-level mixed effects model was employed. Using a random intercept model with PSI conditional on patient, allowed for within patient correlations due to repeated measures on each patient over time. The presence of DCI and sodium nitrite were included as fixed effects, in addition to confounding factors that may influence CBFV dynamics (Age, WFNS score, propofol, P_ET_CO_2_ and noradrenaline infusion levels). All patients were the same Fisher grade. Statistical comparison of the physiological mean scores (baseline vs infusion) was calculated using a two-tailed paired Student’s t-test, after assessing for normality using D’Agostino-Pearson test.

Polar density plots were produced using the Matlab package polarPcolor [51] and represent the percentage time a given phase difference occupied during the recording cycle. The distribution of phase difference at each frequency/scale, enables comparison with stationary phase difference produced using transfer function analysis. A compact display range of 0±45° was used as this represented the range of phase differences that the peaks in PSI occupied.

## 3 Results

### 3.1 Demographics and outcome

10 patients were included in this analysis (mean age 52.3 years; range 44-63 years; 8 Female). All patients were WFNS grade 3-5 on admission and modified Fisher Grade 4. A breakdown of patient characteristics can be found in Table 1. A detailed breakdown of individual sedation and vasopressor requirements during the study can be found in the Supplemental Files.

**Table 1:** Patient Characteristics. L = left; R = right; F = female; M =male; AComm = anterior communicating artery; PComm = posterior communicating artery; A1/A2 = A1 and A2 segment of anterior communicating artery; DCI = delayed cerebral ischaemia; mRS = modified Rankin Scale. Modified Rankin scale (mRS) 0 = no symptoms at all; 1 = no significant disability despite symptoms, able to carry out all usual duties and activities; 2 = slight disability, unable to carry out all previous activities, but able to look after own affairs without assistance; 3 = moderate disability, requiring some help, but able to walk without assistance; 4 = moderately severe disability, unable to walk without assistance and unable to attend to own bodily needs without assistance; 5 = severe disability, bedridden, incontinent and requiring constant nursing care and attention; 6 = dead World Federation of Neurosurgeons (WFNS) grade 1 = Glasgow Coma Scale (GCS) 15, no motor deficit; 2 = GCS 13-14 without deficit; 3 = GCS 13-14 with focal neurological deficit; 4 = GCS 7-12, with or without deficit.; 5 = GCS <7, with or without deficit.

| Patient | Age(year)/<br>Sex | WFNS<br>Grade | Aneurysm<br>Location | DCI | Day DCI<br>Diagnosis | Study<br>Day | mRS at 3<br>Months |
| --- | --- | --- | --- | --- | --- | --- | --- |
| 1 | 55/F | 5 | L PComm | No | n/a | 2 | 1 |
| 2 | 53/F | 5 | A1/A2 | Yes | 10 | 4 | 6 |
| 3 | 52/F | 4 | AComm | No | n/a | 2 | 0 |
| 4 | 50/F | 3 | A1/A2 | Yes | 16 | 3 | 5 |
| 5 | 44/M | 5 | AComm | No | n/a | 4 | 2 |
| 6 | 54/F | 4 | R PComm | No | n/a | 2 | 2 |
| 7 | 56/F | 3 | R PComm | No | n/a | 2 | 1 |
| 8 | 47/F | 4 | L A2 | Yes | 9 | 2 | 2 |
| 9 | 49/F | 3 | R Vertebral | Yes | 8 | 2 | 4 |
| 10 | 63/M | 4 | AComm | No | n/a | 3 | 2 |

The study was performed between days 2 and 4 (mean 2.6) after ictus. At the time of study, all patients were sedated and ventilated and no patient showed clinical or angiographic evidence of DCI or cerebral arterial vasoconstriction. Four (40%) of the patients subsequently developed delayed cerebral infarction. Diagnosis was made by computed tomography scan on the suspicion of the clinical team. One patient died from complications of delayed cerebral ischaemia. Two patients were treated for ventilator acquired pneumonia with IV antibiotics. There were no cases of rebleeding.

### 3.2 Physiological data

There was a significant decrease in mean arterial blood pressure (MAP) following administration of sodium nitrite (mean baseline MAP 77.5 mmHg to mean infusion MAP 71.1 mmHg; p = 0.005, paired two tailed Student’s t test). There were no significant changes in CBFV, ICP, or P_ET_CO_2_ values (Table 2).

**Table 2.** Physiological Data. Mean (standard deviation) ABP = Arterial Blood Pressure; mmHg = Millimetres of Mercury; CBFV = Cerebral Blood Flow Velocity; cm/s = Centimetres per second; etCO2 = End Tidal Carbon Dioxide; kPa = Kilopascals; ICP = Intracranial Pressure

| Patient | ABP (mmHG)<br>p=0.005 |  | CBFV (cm/s)<br>p=0.11 |  | etCO2 (kPa)<br>p=0.53 |  | ICP (mmHg)<br>p=0.77 |  |
| --- | --- | --- | --- | --- | --- | --- | --- | --- |
|  | Pre | During | Pre | During | Pre | During | Pre | During |
| 1 | 68<br>(2.7) | 65<br>(4.3) | 69<br>(8.0) | 69<br>(8.5) | 3.9<br>(0.05) | 4.7<br>(0.02) | nr | nr |
| 2 | 63<br>(3.1) | 61<br>(2.1) | 39<br>(1.8) | 36<br>(2.0) | 3.8<br>(0.04) | 3.5<br>(0.05) | nr | nr |
| 3 | 55 (<br>1.6) | 52<br>(1.1) | 70<br>(2.2) | 66<br>(1.4) | 3.9<br>(0.02) | 3.9<br>(0.03) | 3.3<br>(0.7) | 2.7<br>(0.8) |
| 4 | 77<br>(3.1) | 66<br>(6.1) | 44<br>(2.1) | 40<br>(4.8) | 4.8<br>(0.02) | 4.6<br>(0.02) | 7.5<br>(1.6) | 6.4<br>(3.0) |
| 5 | 55<br>(4.9) | 59<br>(3.4) | 41<br>(3.0) | 47<br>(2.5) | 4.8<br>(0.12) | 4.8<br>(0.12) | 6.6<br>(1.5) | 6.9<br>(1.0) |
| 6 | 72<br>(3.4) | 66<br>(2.6) | 106<br>(5.7) | 98<br>(4.3) | 5.9<br>(0.14) | 5.7<br>(0.11) | 20.4<br>(0.0) | 20.5<br>(0.0) |
| 7 | 88<br>(2.0) | 80<br>(3.4) | 49<br>(3.8) | 49<br>(2.8) | 4.0<br>(0.07) | 4.0<br>(0.08) | 11.4<br>(0.0) | 11.2<br>(0.0) |
| 8 | 84<br>(1.6) | 73<br>(2.0) | 90<br>(7.6) | 91<br>(3.8) | 4.2<br>(0.13) | 4.2<br>(0.06) | 8.5<br>(3.4) | 5.9<br>(3.3) |
| 9 | 100<br>(4.0) | 84<br>(4.6) | 53<br>(2.8) | 47<br>(3.2) | 3.4<br>(0.04) | 3.6<br>(0.08) | 8.9<br>(2.7) | 11.7<br>(3.0) |
| 10 | 113<br>(4.1) | 105<br>(4.1) | 63<br>(4.3) | 60<br>(6.3) | 3.2<br>(0.19) | 3.9<br>(0.06) | nr | nr |

### 3.3 Phase Synchronisation Analysis

Inspection of the group average pre-infusion baseline PSI (Figure 3a) revealed distinct differences when compared to PSI values in healthy volunteers obtained by Latka et al. (Figure 1b).[36] The characteristic peak at 0.33 Hz (scale 15) is present, however, it is absent at 0.1 Hz (scale 50). Moreover, at frequencies < 0.07 Hz (scales > 75) where PSI normally falls rapidly to values below 0.2, it rises to values above 0.4. Inspection of the same group average phase distribution polar density plot (Figure 3b) indicates an in-phase relationship for the phase synchronisation peak at 0.33 Hz. At frequencies between 0.07-0.025 Hz, there is a slightly positive phase difference (CBFV preceding ABP).

Although direct comparisons of the PSI values obtained in the patients included in the study with those derived from healthy volunteers cannot be made, it does enable identification of morphological differences that warrant further examination. Statistical comparison was subsequently performed on mean PSI within two frequency ranges: 0.1316 - 0.0833 Hz to include the broad-based peak at 0.1 Hz and below 0.07 Hz

**Figure 3:**
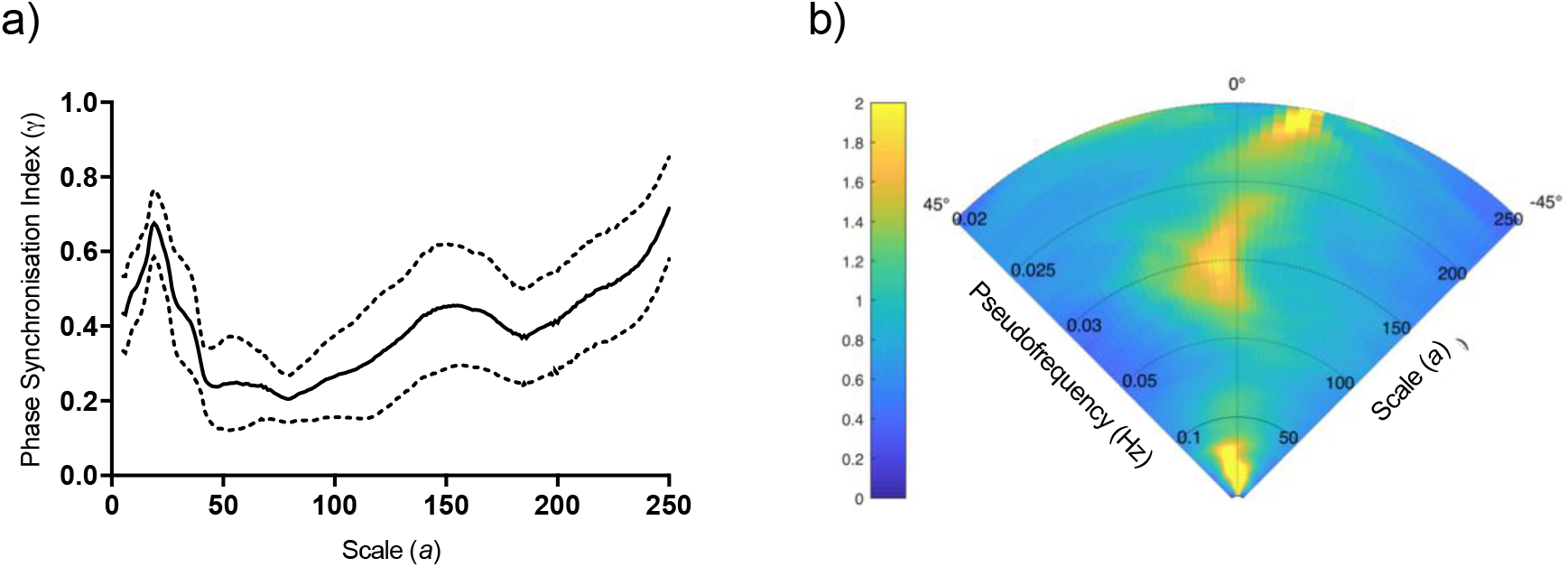
a) Phase Synchronization Index (*γ*) as a function of scale *a* (bottom x-axis) or pseudofrequency *f*_*a*_ (top x-axis) from pre-infusion (baseline) data. Solid line represents the group average, dashed lines correspond to the 95% confidence intervals. b) Polar density plot of group average phase difference distributions over time in pre-infusion (baseline) data. Phase difference is given in degrees, a positive value indicates CBFV leading ABP, whereas a negative value indicates the opposite. Colour bar represents probability of a given phase difference at a given time. The peaks in phase synchronisation observed at 0.33 Hz and 0.07-0.025 Hz, are apparent by their elevated density (and therefore low variance) on the polar plot. The phase difference at scale 0.33 Hz is 0° (in-phase), and slightly positive at frequencies between 0.07-0.025 Hz.

The results of the linear mixed effects modelling of mean PSI for the frequency range encompassing the 0.1 Hz peak are as follows: Within group comparisons revealed that in patients that did not develop DCI, mean PSI significantly increased on the administration of the sodium nitrite infusion (baseline mean PSI = 0.338 (95% CI [0.168, 0.508]) to infusion mean PSI = 0.45 (95% CI [0.285, 0.618]); p value = 0.006). In patients that did develop DCI there was no change in PSI (baseline mean PSI = 0.157 (95% CI [-0.048, 0.363]) to infusion mean PSI = 0.171 (95% CI [-0.033, 0.374]); p value = 0.763).

Within condition comparisons identified that there was no significant difference in mean baseline PSI between patients that developed DCI and those that did not (p value = 0.175). However, there was a significant difference in mean PSI during the infusion (p value = 0.030). These results are illustrated in Figure 4a. Normality and homoscedasticity of the linear mixed effects model was confirmed by examining the qqnorm plot of the model residuals and residuals vs. fitted values for the model respectively (see Supplemental Files). There were no significant effects of Age, WFNS grade, Propofol level, Noradrenaline level and etCO2 level on the mean PSI for the 0.1 Hz peak.

The results of the linear mixed effects modelling of mean PSI for frequencies below 0.07 Hz are as follows: There were no significant within group differences in mean PSI. In patients that did not develop DCI, mean baseline PSI = 0.571 (95% CI [0.421, 0.721]) and mean infusion PSI = 0.601 (95% CI [0.453, 0.748]); (p value = 0.301). In patients that developed DCI, mean baseline PSI = 0.573 (95% CI [0.392, 0.755]) and mean infusion PSI = 0.598 (95% CI [0.418, 0.778]); (p value = 0.458). There were also no significant within condition differences in mean PSI at baseline (p value = 0.984) and during the infusion (p value = 0.982). These results are illustrated in Figure 4b. There was also no significant effect of the additional confounding factors.

**Figure 4:**
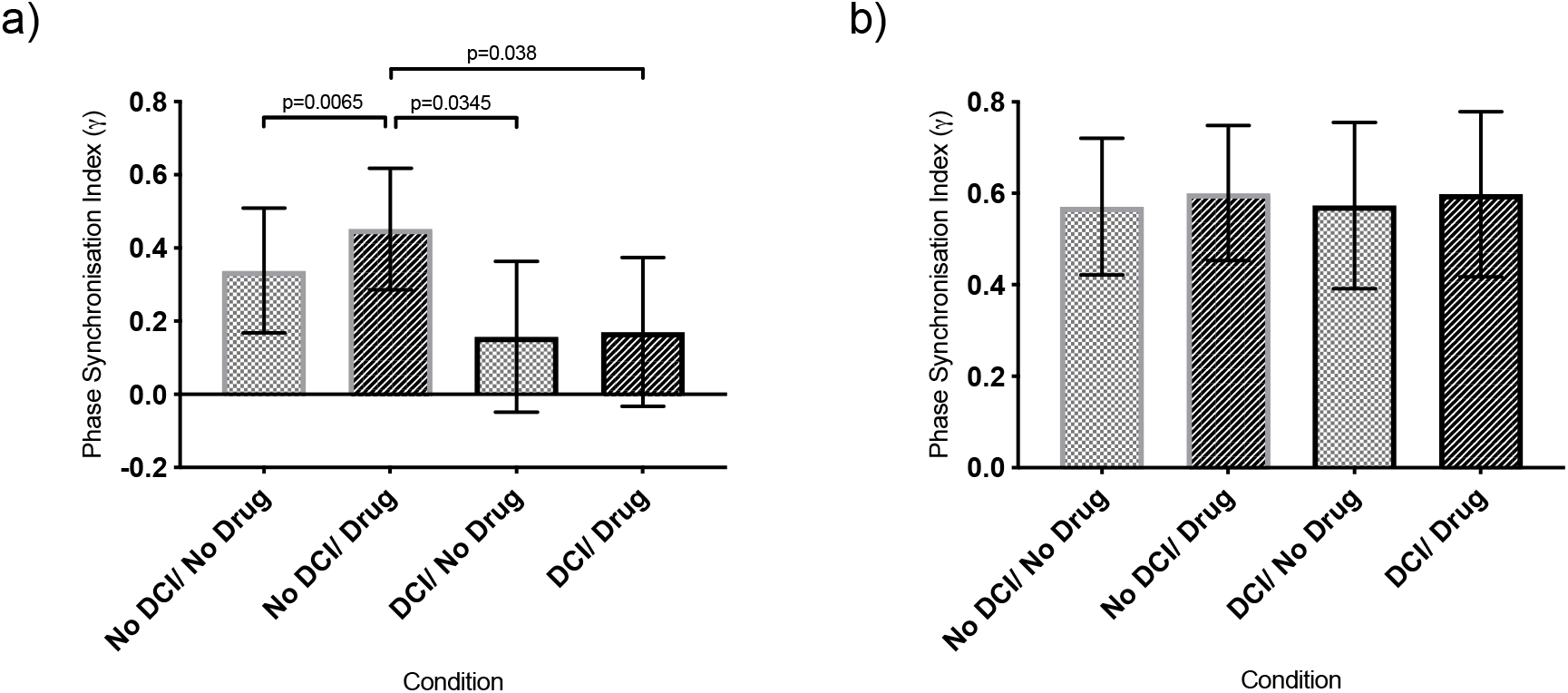
Bar chart of linear mixed effects model estimated mean phase synchronisation index (PSI) (*γ*) for a) for the frequency range encompassing the the 0.1Hz peak and b) for frequencies below 0.07 Hz, for the different conditions. Bars represent 95% confidence intervals. a) A significant increase of PSI in patients that did not develop DCI following sodium nitrite is visible. PSI is also significantly larger in patients that did not develop DCI, than those that did during the infusion of sodium nitrite. b) No significant differences are visible.

Finally, the effect of DCI and sodium nitrite infusion on ABP band power at the frequency range encompassing the 0.1 Hz peak was assessed, to determine if sodium nitrite was inducing a change in ABP power. Band power was taken as the mean absolute-value squared of the wavelet coefficients. Linear mixed effects modelling demonstrated no significant within group or within condition differences.

Figure 5a illustrates the PSI results in a single patient that did not develop DCI, demonstrating a clear increase in PSI around 0.1 Hz following the administration of sodium nitrite. Inspection of the phase distribution polar density plots during the baseline (Figure 5b) and infusion (Figure 5c), show that the increase in PSI around 0.1 Hz has a slight positive phase relationship between CBFV and ABP.

**Figure 5:**
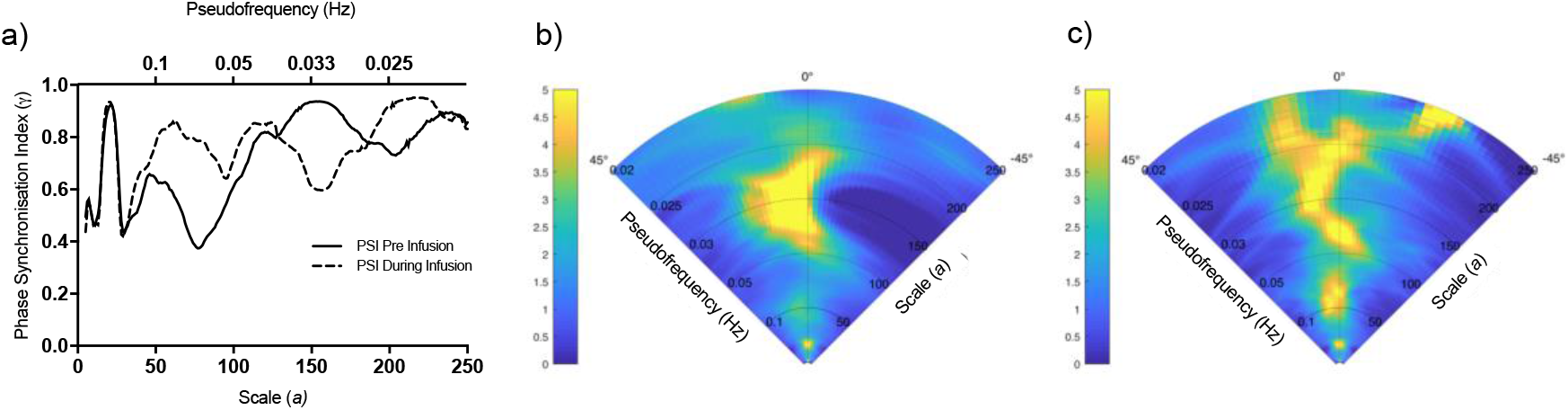
Results from typical patient that did not develop DCI. a) Phase synchronisation index (PSI) (*γ*) before (solid line) and during (dashed line) infusion of sodium nitrite. An increase in PSI around 0.1 Hz is clearly visible. Polar density plots from the same patient before and c) during sodium nitrite infusion. Phase difference is given in degrees, colour bar represents probability of a given phase difference at a given time. An increase in PSI at 0.1 Hz is evident as a corresponding increase in density on the polar plots, with slightly positive phase difference distribution.

## 4 Discussion

### 4.1 Main findings

The aim of this study was to investigate the role of the NO pathway on the phase dynamics of CBF after SAH using an exogenous NO donor. By utilising the continuous wavelet transform and phase synchronisation analysis, we have been able to identify changes in the dynamic phase relationship between CBFV and ABP following the administration of exogenous nitric oxide donor.

Most importantly, we have identified that SAH patients that did not go on to develop DCI demonstrate a significant increase in mean phase synchronisation index in the 0.1 Hz range following the administration of sodium nitrite. Whereas, in patients that did develop DCI, there was no change in mean phase synchronisation index within the same frequency range. Importantly, the examined SAH patients were indistinguishable at baseline with respect to PSI.

### 4.2 Low frequency pulsatile flow

Latka et al. suggested that the peak in phase synchronisation between ABP and CBF at around 0.1 Hz is the result of the synchronisation of low frequency oscillations in arterial blood pressure, known as Mayer waves, and CBF.[36] Mayer waves are thought to result from baroreflex-mediated changes in sympathetic outflow to systemic vasculature [52] and were considered represent an epiphenomenon of normal baroreflex operation with no specific function.[53] Yet a growing body of evidence suggests that they play an essential role in maintaining organ perfusion, especially during periods of physiological stress.

In a study performed by Rickards et al., the physiological responses between subjects exhibiting high and low tolerance to central hypovolemia, were characterised. They demonstrated that the linear coupling between the low frequency oscillations in MAP and CBF, as assessed by spectral coherence, increased progressively with increasing levels of lower body negative pressure in the high tolerance group, but did not change in the low tolerance group. This was despite similar reductions in absolute ABP and CBF between groups.[54] These findings were expanded upon by Romero et al. who demonstrated a greater increase in ABP and CBF low frequency coherence during head-up tit and dehydration over euvolemia.[55] Subsequently, Lucas et al. demonstrated that, by increasing ABP low frequency power, using fixed respiration at 6 breaths per min (0.1 Hz), there was a significant increase in tolerance to combined head-up tilt and lower body negative pressure, when compared to normal breathing. Again, the reduction in CBF was similar between the two breathing patterns.[56] Finally, Anderson et al. recently demonstrated an attenuation of the decrease in cerebral oxygenation, as assessed by near infrared spectroscopy, and increased tolerance to lower body negative pressure during induced pressure oscillations at 0.1 Hz when compared to lower body negative pressure with no oscillatory variations.[57]

Spectral coherence estimates the linear phase-consistency between frequency decomposed signals and is an estimate of the coupling between oscillatory signals, akin to phase synchronisation. These studies, therefore, demonstrate that increased tolerance to central hypovolemia is associated with the coupled oscillatory pattern of ABP and CBF around 0.1 Hz. This process has been termed low frequency pulsatile flow.[58] As previously discussed, Garry et al, demonstrated an increase in the EEG alpha delta ratio[24], suggestive of improved CBF.[25] However, the primary analysis conducted by Garry et al. identified no change in CBF as assessed by time averaged CBFV measures. It has been proposed that rhythmic fluctuations in ABP provide a pump-like effect improving tissue perfusion during physiological stress via increases in the pressure gradient within the vasculature.[59] Alternatively, there is evidence from computational models to suggest that oscillatory flow is beneficial to radial oxygen diffusivity.[60]

Although a direct comparison with the PSI in healthy volunteers produced by Latka et al.[36] is subject to several caveats (see limitations), the apparent absence of the peak in PSI around 0.1 Hz in the baseline data is suggestive of impaired baseline cerebral perfusion. Unfortunately, comparison with previous studies investigating CBF control in SAH is difficult. The majority of studies utilised linear correlation at frequencies < 0.05 Hz as a metric of CBF control[6–9,14], and did not report results on the coupling between ABP at CBF around 0.1 Hz. In the single study that performed a frequency domain analysis[11], the reported frequency band (0.03 Hz - 0.15 Hz) encompasses much of the very low frequency domain (0.02-0.07 Hz).[29] As will be expanded upon later, an increased coherence in the very low frequency band is associated with impaired dynamic cerebral autoregulation, as would be likely after SAH, and would therefore mask any reduction in coherence we may expect to observe around 0.1 Hz.

Despite this, a comparable study using continuous wavelet transforms and PSI in traumatic brain injury patients also identified the absence of the 0.1 Hz centred peak in PSI when compared to matched controls.[40] This suggests that disruption to low frequency pulsatile flow may represent a common pathophysiological process in acute brain injuries.

### 4.3 Role of Nitric Oxide

The increase of 0.1 Hz-centred phase synchronisation following administration of sodium nitrite suggests that NO plays a critical role in the synchronisation between low frequency oscillations in ABP and CBF. Furthermore, the absence of the 0.1 Hz centred peak in the baseline synchronisation index supports evidence suggesting functional disruption to the NO signalling pathways during EBI.[15]

There is a strong biological plausibility to the above assertion. Pulsatile flow is known to induce endothelial sheer stress and the liberation of NO[61], which in turn induces vascular smooth muscle relaxation via cyclic guanosine monophosphate (cGMP) dependent processes.[62] Vasodilatation of peripheral resistance vessels downstream of the insonated middle cerebral artery segment would promote flow and therefore increase CBFV in the insonated segment.[63] Furthermore, inhibition of NO synthase in isolated cerebral blood vessels has been shown to induce chaotic vasomotion at 0.1 Hz[64] and such decoupled behaviour would also result in impaired synchronisation. Finally, Nafz et al. demonstrated that superimposed 0.1Hz oscillations improved renal excretory function, in the context of reduced renal perfusion pressure, via increased generation of NO.[65] Increasing cerebral NO, through sodium nitrite administration, may restore NO levels to an effective concentration during periods of functional NO depletion.

However, evidence in healthy human volunteers is far from conclusive. Using the non-specific NO synthase inhibitor NG-monomethyl-L arginine (L-NMMA), Zhang et al. failed to demonstrate an effect on low frequency coherence during head-up tilt, despite inducing a similar increase as previously described.[32] Furthermore, Lavi et al. were unable to demonstrate that the NO donor, sodium nitroprusside (SNP), modulated mechanoregulation by assessing changes in CBF to gradual increases in ABP using phenylephrine.[30] However, White et al., using L-NMMA and the thigh cuff test, demonstrated a significant decrease in the autoregulation index, signifying impaired autoregulation when compared to controls with matched blood pressure changes.[31]

The findings of Lavi et al. are not surprising, derived absolute changes in CBF from transcranial Doppler recordings may be inaccurate if changes in vessel diameter occur, which may be expected with SNP. Unfortunately, the autoregulation index used by White et al. is a single value given to represent a number of fitted parameters. As such, it is difficult to directly compare with our results. It is worth pointing out that the dose of L-NMMA used by White et al. was over double that used by Zhang et al. and, in combination with uncertainty of the degree and the specificity of L-NMMA inhibition of NOS activity in the cerebral circulation, may in part explain the differential findings.

However, the discordance between the findings of Zhang et al., which is the most directly comparable study, and our own are clear. It is possible to reconcile these differences by considering the indirect action of NO. In addition to the direct action on vascular smooth muscle cells, NO also inhibits the synthesis of the vasoconstricting 20-hydroxyeicosatetraenoic acid (20-HETE).[66] This effect may underlie a significant fraction of the dilating effect of NO.[67] SAH is associated with increased levels of 20-HETE[68] which may provide a mechanism to explain the differential effect of NO in health and disease. This is further supported by the observation that the sodium nitrite administration in healthy volunteers does not influence EEG alpha/delta ratios.[69]

### 4.4 Differential Outcome Response

Our findings clearly demonstrate that the increase in 0.1 Hz centred phase synchronisation following administration of sodium nitrite occurs in patients that did not go on to develop DCI but was not present in those patients that did develop DCI. These findings replicate the differential EEG response also observed by Garry et al.[24] A possible explanation for the differential response is that the patients who developed DCI sustained a more severe injury in the acute period following aneurysmal rupture with a resulting greater functional NO depletion. Furthermore, there is evidence to suggest a correlation between subarachnoid haemorrhage blood volume and the development of DCI.[70,71] Elevated free haemoglobin and its breakdown products would result in increased buffering of both endogenous NO and NO liberated from the infused sodium nitrite.

If the result of decreased NO bioavailability in those who develop DCI, we may expect to observe an increase in phase synchronisation with longer or larger infusions. However, it is likely that changes in downstream signalling factors also contribute significantly. Animal models have shown both reduced expression of vascular smooth muscle soluble guanylate cyclase[72] and increased rates of cGMP hydrolysis[73] during DCI. Additionally, higher levels of 20-HETE in the CSF of patients has been associated with the development of DCI.[68]

Modelling work by Catherall showed that cGMP only has influence over vascular smooth muscle phosphorylation, and, therefore, contractility, over a limited range of intracellular calcium concentrations.[74] Excitotoxicity, resulting in elevated intracellular calcium concentration, is a known pathophysiological component of SAH.[75] It is plausible that in patients that develop DCI, modulating cGMP concentrations via NO will have no effect due to significantly elevated intracellular calcium concentrations. This correlates with evidence that the calcium channel antagonist, nimodipine, improves outcomes following SAH.[76]

Alternatively, functional NO depletion maybe a characteristic of EBI only in patients that do not develop DCI. Whereas, in patients that develop DCI cerebral NO concentration, via iNOS dependent production, have exceeded the functional operating range, potentially reaching cytotoxic levels, rending additional increases ineffectual. This would be consistent with evidence demonstrating higher NO breakdown products in patients that develop DCI [20,21], and the hypothesis that NO mediated toxicity results in DCI.[15]

### 4.5 Dynamic Cerebral Autoregulation

When considered in the frequency domain, normal dCA is characterised by low coherence, low gain and high phase lead in the very low frequency range, and increasing coherence and gain, and decreasing phase lead as frequencies increase.[29]

Applying this interpretation, the high coherence observed at 0.1 Hz during central hypovolemia should be considered to reflect impaired dCA. However, as previously described, the increase in coherence appears to protect cerebral perfusion, suggesting a relationship more complex than that of a simple high pass filter. Given that the majority of studies investigating dCA in SAH have only considered the very low frequency band (0.02 - 0.07 Hz) and the apparent complex relationship at higher frequencies, we will regard frequencies below 0.07 Hz to reflect dCA.

Latka et al. considered a decreased phase synchronisation at frequencies below 0.07 Hz to be reflective of intact dCA.[36] Although Mitsis et al., using Volterra kernel analysis, and Peng et al., using wavelet phase synchronisation, have both attributed a proportion of the low phase synchronisation below 0.07 Hz to CO2 variability (see limitations), it was still characterised by a fall at 0.07 Hz.[33,41] While subject to the same caveats as before, the apparent increase in baseline phase synchronisation at frequencies below 0.07 Hz would appear to indicate impaired dCA. This is further exemplified by the reduced phase lead, suggesting passive flow rather than dynamic counter regulation. These findings are in line with other dCA studies in SAH which, although predominantly performed using time domain linear correlation, show a stronger linear relationship between ABP-CBFV at frequencies below 0.05 Hz. [6–9,14]

Using linear correlation, Budohoski et al. demonstrated that disturbed autoregulation in the 5 days after SAH was associated with DCI at 21 days.[6] We failed to demonstrate a difference in baseline phase synchronisation below 0.07 Hz between patients that developed DCI and those that did not. This is likely a reflection of the population size rather than the analysis technique (98 patients were recruited into the study by Budohoski et al.). We also failed to demonstrate an effect of sodium nitrite on phase synchronisation below 0.07 Hz in either group. This may suggest that disruption to NO pathways does not play a role in very low frequency dCA. However, Tseng et al. demonstrated a reduced duration of impaired dCA in patients treated with pravastatin.[77] Statins have been proposed to recouple endothelial NO synthase function, resulting in increased NO levels.[78] One explanation for the failure to observe a significant difference may be the reduced length of data available at low frequencies. As frequencies decrease the COI increases, reducing the available data for analysis.

### 4.6 Limitations

The small sample size and absence of a control group represent the major limitations of the present study. Furthermore, we have used morphological differences in the PSI between our patient population and healthy volunteers, derived from Latka et al.,[36] to guide subsequent analysis. However, caution must be applied in drawing conclusions regarding the aetiology of these differences. Our patient population was sedated and mechanically ventilated, with many of them receiving vasopressor drugs. These factors alone may result in significantly different phase synchronisation characteristics and, although attempts have been made to account for within patient differences, they limit the ability to draw firm comparisons. Future studies should include a control population drawn from sedated and ventilated patients without acute neurological injury.

Unmeasured variability, in the form of natural oscillations of P_ET_CO_2_, may account for a proportion of the low phase synchronisation at certain frequencies.[33,41] However, these studies was performed in spontaneously breathing healthy volunteers. Given that in our study all the patients were mechanically ventilated with pressure controlled intermittent positive pressure ventilation, the variability of P_ET_CO_2_ is very low. As such, there appeared no advantage to adding additional complexity by incorporating P_ET_CO_2_ into the analysis approach. Furthermore, inclusion of P_ET_CO_2_ had no effect on PSI estimates at 0.1 Hz and therefore would be unlikely to influence our primary finding. Average P_ET_CO_2_ levels were, however, considered in the mixed effects linear model.

### 4.7 Conclusion

We have demonstrated that pharmacological manipulation of *in vivo* NO levels in patients with high grade SAH results in increased synchronisation between ABP and CBF around 0.1 Hz in patients that do not develop DCI. Such synchronisation has been proposed as a mechanism by which cerebral perfusion is maintained during periods of physiological stress. These findings suggest that functional NO depletion plays a role in impaired CBF control during EBI, but that the development of DCI may have a distinct pathophysiological aetiology.

## Abbreviations

20-HETE: 20-Hydroxyeicosatetraenoic Acid
ABP: Arterial Blood Pressure
CARNET: Cerebral Autoregulation Network
CBF: Cerebra Blood Flow
CBFV: Cerebral Blood Flow Velocity
cGMP: Cyclic Guanosine Monophosphate
COI: Cone of Influence
CSF: Cerebrospinal Fluid
dCA: Dynamic Cerebral Autoregulation
DCI: Delayed Cerebral Ischemia
EBI: Early Brain Injury
EEG: Electroencephalogram
ICP: Intracranial Pressure
L-NMMA: NG-monomethyl-L arginine
MAP: Mean Arterial Pressure
MCA: Middle Cerebral Artery
mRS: Modified Rankin Scale
PETCO2: End Tidal CO2
PETO2: End Tidal O2
PSI: Phase Synchronisation Index
SAH: Subarachnoid Haemorrhage
SNP: Sodium Nitroprusside
TCD: Transcranial Doppler Ultrasonography
WFNS: World Federation of Neurosurgeons

## Supplemental Files

**Figure 1:**
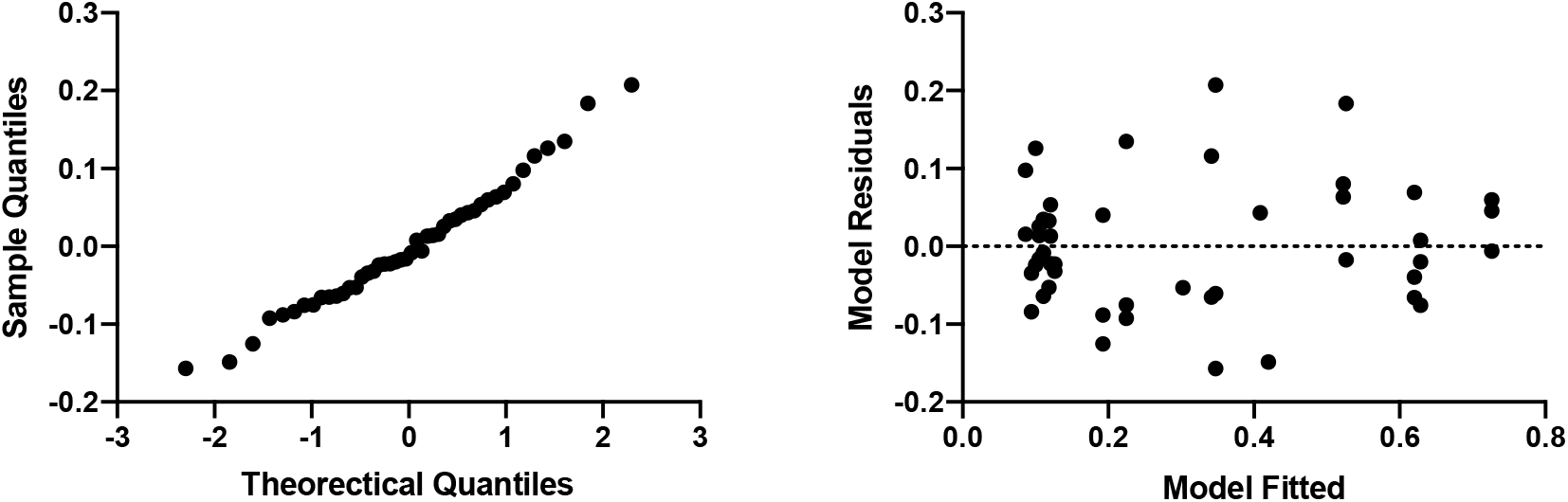
Diagnostic plots to show the a) normality of the residuals in the linear mixed effects model and b) the fitted model is unbiased and homoscedastic. These plots demonstrate that the model is a good fit for the data.

**Table 1:** Sedation, paralysis and Inotropes. mg/hr = milligrams per hour; mcg/hr = micrograms per hour; mcg/kg/min = micrograms per kilogram per minute

| Patient | Propofol (mg/hr) |  | Fentanyl (mcg/h) |  | Midazolam (mg/hr) |  | Atracurium (mg/hr) |  | Noradrenaline (mcg/kg/min) |  |
| --- | --- | --- | --- | --- | --- | --- | --- | --- | --- | --- |
|  | Pre | During | Pre | During | Pre | During | Pre | During | Pre | During |
| 1 | 0 | 0 | 200 | 200 | 10 | 10 | 0 | 0 | 0.34 | 0.34 |
| 2 | 0 | 0 | 300 | 300 | 20 | 20 | 50 | 50 | 0.38 | 0.38 |
| 3 | 200 | 200 | 150 | 150 | 0 | 0 | 0 | 0 | 0 | 0 |
| 4 | 180 | 200 | 0 | 0 | 0 | 0 | 0 | 0 | 0.17 | 0.2 |
| 5 | 100 | 100 | 150 | 150 | 0 | 0 | 0 | 0 | 0.15 | 0.15 |
| 6 | 0 | 0 | 100 | 100 | 0 | 0 | 0 | 0 | 0.14 | 0.14 |
| 7 | 220 | 230 | 200 | 250 | 0 | 0 | 0 | 0 | 0.25 | 0.25 |
| 8 | 150 | 150 | 100 | 100 | 0 | 0 | 0 | 0 | 0.15 | 0.16 |
| 9 | 180 | 180 | 200 | 200 | 0 | 0 | 0 | 0 | 0.2 | 0.2 |
| 10 | 150 | 200 | 200 | 200 | 0 | 0 | 0 | 0 | 0 | 0 |

